# cis dominantly explains regulatory divergence between two indica rice genotypes; drought further enhances regulatory differences

**DOI:** 10.1101/714907

**Authors:** Nelzo C. Ereful, Antonio Laurena, Li-Yu Liu, Shu-Min Kao, Eric Tsai, Michael Thomson, Andy Greenland, Wayne Powell, Ian Mackay, Hei Leung

## Abstract

**Abstract** cis and/or trans regulatory divergence within or between related taxa on a genome-wide scale has been largely unexamined in crops, more so, the effect of stress on cis/trans architecture. In this study, the indica genotypes IR64, an elite drought-susceptible lowland variety, and Apo (IR55423-01 or NSIC RC9), a moderate drought-tolerant upland genotype together with their hybrid (IR64 × Apo) were exposed to non- and water-stress conditions. Evidence of cis and/or trans regulatory differences was tested between these two indica rice genotypes. By sequencing (RNA-seq) the parents and their hybrid, we are able to map genes diverging in cis and/or trans factors between the two genotypes. Under non-stress conditions, cis dominantly explains (11.2%) regulatory differences, followed by trans (8.9%). Further analysis showed that water-limiting conditions largely affect trans and cis + trans factors. Between the two inbred lines, Apo appears to exhibit higher expression fold change of genes enriched in “response to stress” and “photosynthesis” under non- and water-stress conditions. On the molecular level, cis and/or trans regulatory divergence explains their genotypic differences and differential drought response. Parent–hybrid RNA-seq has the potential to identify genes diverging in cis and/or trans factors even between intra-sub-specifically related genotypes. By comparing cis/trans landscape under stressed and unstressed conditions, this approach has the ability to assess the impact of drought on gene expression. Computational analysis and association of several drought-yield QTL markers with cis-diverging genes provide converging evidences suggestive of a potential approach to identify trait-associated candidate genes using hybrids and their parents alone.

**Key Message:** cis dominantly explains divergence of two indica rice genotypes, IR64 and Apo under normal conditions while trans and cis + trans regulatory factors are largely affected by drought

## Introduction

Allele-specific expression (ASE) imbalance or simply allelic imbalance (AI) happens when one of the two alleles in a heterozygote is expressed more than the other. It has been a subject of investigations in several organisms including crops (rice, barley and maize), model animals (e.g. mouse, *Arabidopsis, Drosophila*) and humans. In a hybrid, both alleles are exposed to the same nuclear environment, thus asymmetric expression of the hybrid alleles is attributed to cis regulatory polymorphism. On the other hand, total expression differences between two genotypes which cannot be explained by cis-regulatory differences is ascribed to trans (Wittkopp et al. 2004; McManus et al. 2010). Hence, expression differences between the parental genes that disappear in the hybrid are caused by trans effects (Tirosh et al. 2009).

The relative contribution of both regulatory and coding sequences in generating phenotypic variation and thus evolutionary differences and distinct ecological adaptation has been a subject of recent reports. Regulatory sequences, in particular, have been associated with evolutionary and adaptive divergence in humans (Buil et al. 2015), *Drosophila* (Wittkopp et al. 2004; Wittkopp et al. 2008; McManus et al. 2010; Gompel et al. 2005), yeast (Tirosh et al. 2009), *Arabidopsis* (He et al. 2016), tomato (Guerrero et al. 2016) and stickleback (Verta and Jones, 2019). Some of these reports reveal that it is the changes in cis-acting regulatory sequences that contribute predominantly to phenotypic divergence (Carroll et al. 2008; Stern et al. 2008; Wittkopp et al. 2008; He et al. 2016). However, other investigations have claimed that there is an equal contribution of both cis and trans regulatory factors to evolutionary divergence (Schmitz et al. 2016; Guerrero et al. 2016).

Despite the increasing information of the participation of local and distal regulatory sequences to the expression divergence of organisms, regulatory differences in rice, arguably the world’s most important crop, is poorly understood. Moreover, little is known on the effect of stress as a selective agent on gene regulation. A report on rice demonstrated that regulatory divergence is associated with ecological speciation and adaptive alterations (Guo et al. 2015).

A 2015 study shows that modern Green Revolution (GR) rice varieties are typically drought-sensitive due to tight linkage between the loci involved in drought tolerance and plant height (Vikram et al. 2015). Released 50 years ago, IR8, the GR rice is a high-yielding semi-dwarf rice variety, and is derived from a cross between Peta and Dee-geo-woo-gen (DGWG), both of which are *indica* (Hargrove and Coffman, 2006). With a coefficient of parentage (COP) of 0.131 (see pedigrees, Supplementary Information, S1), IR64 and Apo, the materials used in this study, have descended from IR8. Despite their common ancestry, these genotypes show differential response to drought conditions and contrasting ecological adaptation – IR64 thrives in a lowland ecosystem; Apo, in upland. Understanding their regulatory differences may therefore provide insights on the molecular basis of their varying drought response and contrasting adaptation.

Majority of previous studies explored regulatory divergence between or within two or more interspecifically related organisms (e.g., *Drosophila, Arabidopsis*, yeast). In this study, two intra-sub-specifically related rice genotypes (both indica) were tested for cis- and trans-regulatory differences on a genome-wide scale. This may shed hints on the resolving power of mRNA-seq to dissect regulatory divergence between highly genetically related inbred lines. Such strategy may enable the identification of loci with divergent regulatory regions and potentially causative polymorphism between closely related genotypes.

Our investigation is limited to one-way specific cross (IR64 × Apo) to provide preliminary information on the regulatory differences between the two rice genotypes grown under normal (well-watered) and water-stress conditions. Such environmental perturbation may further drive regulatory differences.

## Results and Discussion

The genotypes IR64, Apo and their hybrid (IR64 × Apo) were exposed to both non- (control) and water-stress conditions at early flowering stage and were sequenced using paired-end mRNA-seq (3 genotypes × 2 treatments × 2 replicates) (see Materials and Method). We then tested these datasets for evidence of cis and/or trans regulatory differences. Intra-sub-specific comparison of two rice genotypes with contrasting tolerance against water-limiting conditions allows the detection of regulatory differences at candidate loci putatively involved in their distinct ecological adaptation and divergent response against drought. In this paper, “cis and/or trans” means cis, trans, and their interactions which include cis + trans (synergistic), cis × trans (antagonistic), and compensatory.

To be consistent with our previous papers (Ereful et al. 2016; Ereful et al. 2020) and allow comparison of expression data, we used the *O. sativa* ssp. *japonica* (cv. Nipponbare) MSUv7 cDNA to map the parental reads. Variant calls (SNPs and Indels) between the two inbred parental lines were identified to discern which parent-specific allele belongs to which parental genotype in the F1. We then created a pseudo-reference to map the heterozygote reads and analyzed for cis/trans regulatory divergence. In the hybrid, since both genotype-specific alleles are exposed to the same trans-acting factors, asymmetric expression is attributed to cis differences or allele-specific epigenetic changes (Cowles et al. 2002; Wittkopp et al. 2008).

Using this approach, 9141 genes were found to have one or more mapped reads across genotype–line–treatment combinations (Table S1). Only genes with a total parental read count of at least 20 in each treatment were considered for further analysis (IR64 + Apo ≥ 20). We computed the expression ratios (log_2_-transformed) of genes between the two parental genotypes (Parental, P = log_2_(IR64/Apo)) and between the two parent-specific alleles in the F1 heterozygotes (Cis, C = log_2_(IR64_F1_/Apo_F1_)) (see Materials and Methods); trans differences are computed as, T = P – C. By plotting the log_2_-transformed expression ratios of the parents (P) against the hybrid (C) we were able to map isoforms diverging in cis and/or trans regulatory factors and estimate their contributions to the total divergence of the two genotypes under non- and water-stress conditions.

If cis-regulatory differences fully explain divergence between the two genotypes, points (i.e. isoforms) will lie closely on the y=x curve (tested by binomial/Fisher’s exact tests, FDR < 0.5%; see Materials and Method for complete statistical tests). On the other hand if trans-regulatory divergence explains their differences, points (i.e. isoforms) will lie at y=0 curve (exact tests, FDR <0.5%) (Wittkopp et al. 2004; Wittkopp et al. 2008; McManus et al. 2010). Additionally, cis + trans (synergistic), cis × trans (antagonistic), and compensatory effects may also explain their differences. Genes exhibiting cis × trans difference lies closely on the y = –x curve; compensating on the x = 0 curve; for cis + trans, see Fig. 1.

**Fig. 1.**
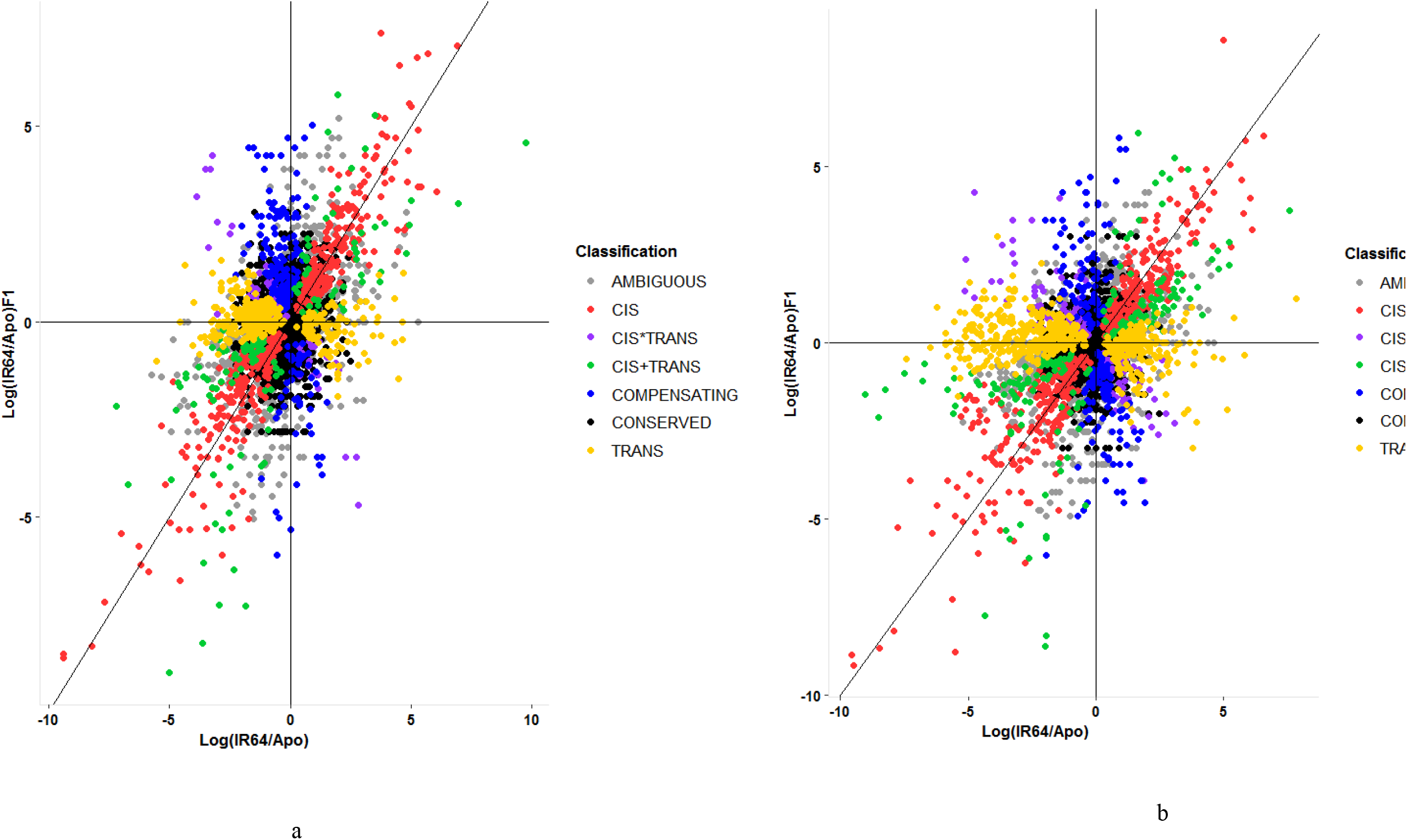
The “scattering” effect of drought stress on cis and/or trans regulatory architecture between two indica genotypes: IR64 and Apo under (a) non- and (bB) water-stress conditions. Visual inspection of the two graphs shows more dispersed points of B compared to A. X-axis indicates expression ratios (in log_2_) between the parental genotypes, Log_2_(IR64/Apo); Y-axis, between the two parent-specific alleles in the hybrid, Log_2_(IR64/Apo)F1.

### Regulatory divergence under non-stress conditions

Under non-stress conditions, 3705 of the 9141 isoforms were found to have a minimum total mapped read of 20 or more between the parents (IR64 + Apo ≥ 20). Of these, 415 (11.2%) and 328 (8.9%) of the 3705 isoforms exhibit cis- and trans-regulatory divergence, respectively, between the parents (Table 1; Fig. 1; see Table S2 for complete list of isoforms diverging in cis and/or trans regulatory factors under non-stress treatment). Apparently, cis dominantly explains the regulatory differences over the other categories between IR64 and Apo (exact tests, FDR<0.5%). (“Conserved” and “ambiguous” categories were omitted from this comparison). These are shown in red points (i.e. isoforms) lying within or close to the y=x curve (Fig. 1A). GO enrichment analysis of isoforms diverging in cis using AgriGO (Tian et al. 2017) showed significant enrichment of genes associated with “response to stress” (biological process, FDR<0.05; 70 genes; Fig. S1) and “nucleolus” (cellular component, FDR<0.05; 15 genes; Fig. S2). The former suggests that their contrasting response against stress is explained by cis divergence. There is now a growing evidence of the participation of nucleolus to abiotic stress including drought (reviewed in Kalinina et al. 2018).

**Table 1.** Summary of the number of isoforms exhibiting cis and/or trans regulatory divergence between IR64 and Apo under non- and water-stress conditions. χ^2^ statistics (with P-values; test for 2×2 contingency or two independent proportions with Yates’ continuity correction) indicates the significant difference of the number of isoforms classified in each regulatory category between the two treatments.

| Regulatory factors/interactions | Non-stress<br>No. of genes (%) | Water-stress<br>No. of genes (%) | $\chi^2$ statistics<br>(P-value) |
| --- | --- | --- | --- |
| cis | 415 (11.2) | 456 (12.6) | 3.19 (0.074) |
| cis + trans<br>(synergistic) | 75 (2.0) | 194 (5.4) | 56.45 (< 0.001) |
| cis × trans | 135 (3.6) | 118 (3.3) | 0.72 (0.397) |
| compensatory | 187 (5.1) | 188 (5.2) | 0.05 (0.828) |
| trans | 328 (8.9) | 551 (15.2) | 69.34 (< 0.001) |
| Ambiguous | 696 (18.8) | 752 (20.7) | 4.315 (0.038) |
| Conserved | 1869 (50.4) | 1366 (37.7) | 120.529 (< 0.001) |
| Total | 3705 | 3626 |  |

On the other hand, GO analysis of isoforms diverging in trans under non-stress conditions showed significant enrichment of genes associated with “photosynthesis” and “generation of precursor metabolites and energy” (biological process, FDR<0.05; Fig. S3) and “thylakoid” and “plastid” (cellular components, FDR<0.05; Fig. S4). Analysis of the log_2_-transformed expression ratios between the two parental genotypes showed that Apo exhibited higher expression fold change (FC) over IR64 in all of the 21 genes enriched in photosynthesis, 18 of which showed at least 2× FC. This photosynthetic efficiency of Apo in terms of expression is ascribed to trans and may further account for its tolerance against drought.

Collectively, cis and trans largely explain divergence of the two genotypes, although compensatory (5.1%), cis + trans (synergistic, 2.0%) and cis × trans (antagonistic, 3.6%) also contribute to these differences, albeit modestly. GO enrichment analysis of genes diverging in cis + trans showed significant association with “response to stress” and “response to stimulus” (biological process; FDR<0.05) most of which were significantly expressed in the parent Apo (76% or 22 of the 29 genes significantly expressed at FC≥2×). Interestingly, one of the genes diverging in cis + trans is the transcription factor ‘no apical meristem’ (LOC_Os12g29330), identified to be a mainstay in the drought-response region in the Vandana/Way Rarem cross (OsNAM_12.1_; Dixit et al. 2015).

#### Cis/trans architecture largely explains genotypic differences

Overall differences between the two parental genotypes were determined using DESeq2 (Love et al. 2014), genotypic differential expression or GDE, to capture loci varying between IR64 and Apo across the treatments. Using the 9141 isoforms (described above), GDE between the parents showed 146 genotypically differentially expressed genes (at FC≥2 or |log_2_FC|≥1, FDR<0.05; Table S3). Of these GDE genes, 89 (or 61%) exhibited cis divergence between the genotypes; 40, cis + trans (27.3%); 9, trans (6%). Taking all these proportions together, 94% (or 138 of the 146) of the genotypic variations are explained by cis and/or trans regulatory differences between the genotypes. Hence, on the molecular level, cis/trans regulatory landscape explains GDE.

Apparently, local regulatory variations explain mostly of the genotypic differences. Reports showed that natural selection reduces the level of variability of local genomic regions leaving molecular patterns of selection such as increased genetic divergence around adaptive loci (Nielsen et al. 2005; Verta and Jones, 2019; Wachowiak et al. 2018). Modern selection and recent breeding procedures have likely generated these differences in which most cis/trans regulatory factors and their interactions appear to favor Apo. These features possibly explain their varying ecological adaptations to lowland (IR64) and upland (Apo) ecosystems and the susceptibility of IR64 and tolerance of Apo to water-limiting conditions.

Our finding is consistent with previous reports which suggests that cis-regulatory changes explain adaptive and ecological divergence between two species of rice (*O. rufipogon* and *O. nivara*; Guo et al. 2015) and among species of *Arabidopsis* (*A. lyrata, A. halleri* and *A. thaliana*; He et al. 2016). Furthermore, the finding that cis effects can explain interspecies divergence is common in nature as was previously reported between two subspecies of rice (indica and japonica; Campbell et al. 2019), in *Drosophila* (Wittkopp et al. 2004; Wittkopp et al. 2008; Osada et al. 2017), coffee (Combes et al. 2015), and most recently stickleback (Verta and Jones, 2019).

### Regulatory differences under water-limiting conditions

To examine how water-stress treatment changes the regulatory landscape, all lines (parental and hybrid) were exposed to water-limiting conditions. Using the same pipeline as the non-stress treatment, 3626 of the 9141 isoforms were found to have a minimum total mapped read of 20 or more between the parents (IR64 + Apo ≥ 20) under water-stress conditions. Their cis/trans regulatory landscape under drought stress is plotted in Fig. 1B. Initial comparative visualization of the two figures (Fig. 1A vs B) reflects the impact of stress as a mechanism to propel cis/trans regulatory differences. Such change in the molecular landscape is suggestive of the effect of drought as an agent of selection for gene regulation.

Of the 3626 isoforms, 456 (or 12.6% of the 3626) and 551 (15.2%) of genes exhibit cis- and trans-regulatory divergence, respectively, which could explain differences between the two genotypes under stress conditions (Table 1; Fig. 1B; see Table S4 for complete list of genes). These results suggest that under water-stress conditions, trans-acting regulatory variations dominantly explain their differences, superseding cis. The χ^2^ test confirms that it is the trans and cis+ trans that are significantly affected by the abiotic stress (Table 1), although trans is more significantly affected (χ^2^=69.34) than the latter (χ^2^=56.45; P<0.001). Similar to a previous finding in *Arabidopsis* (Cubillos et al. 2014), cis is relatively stable under water perturbations (χ^2^=3.19; P=0.074).

The finding that trans regulation differs between environmental regimes agrees with previous studies in yeast (Tirosh et al. 2009) and *Arabidopsis* (Cubillos et al. 2014). We suspect that, as some trans factors are mobile elements (expression; e.g. transcription factors and non-coding RNAs) and can diffuse outside the nucleus, they have been primarily affected by the abiotic stress. Proteins such as TFs have been known to be directly exposed to evolutionary and selection pressures. Additionally, the sensitivity of trans to changes in environmental conditions may be ascribed to its broader mutational target size (sequence) compared to cis mutations (Wittkopp et al. 2004; Metzger et al. 2016; Metzger et al. 2017). Similar to non-stress conditions, trans effects under drought conditions showed GO enrichment in “photosynthesis” (Fig. S5), “thylakoid” and “plastid” (Fig. S6). Apo demonstrates a higher expression fold change over IR64 across all genes enriched in “photosynthesis” (FC≥2, FDR<0.05; 20 genes). Apo appears to photosynthesize more efficiently even under water-limiting regime and is ascribed to trans like it is under non-stress conditions.

#### cis/trans architecture explains differential drought response

Using 3-way DE (referred here as drought DE, DDE), expression variations that arise due to the interactions between the parental genotypes and environment can be detected (G × E) (see Materials and Methods). Results showed that 32 isoforms respond to drought stress across genotypes (Table S5), 13 of which exhibit trans regulatory differences; 12, cis + trans; 3, cis and; 2, cis × trans. This means that 94% (or 30 of the 32 isoforms) of the drought-responsive genes is explained by cis/trans architecture under water stress. Hence, on the molecular level, the two genotypes’ varying response against drought is explained by cis/trans regulatory differences.

### Test of ASE in the hybrid

In the previous test above, we showed the cis and/or trans landscape of the two parental genotypes and their hybrid under two contrasting conditions. As allelic imbalance is ascribed to cis differences in a heterozygote state, we further assess the effect of water-limiting conditions on asymmetric expression independent of the parents. This entails plotting the calculated log_2_-transformed expression ratios of the genotype-specific alleles of the hybrid under non-stress or control (x-axis) and water-stress (y-axis) (with binomial exact test at FDR<0.5%, against the null hypothesis of having no ASE). This gives information on which isoforms exhibit allelic imbalance under non- and/or water-stress conditions in the hybrid. Results are depicted in Fig. 2 and are summarized in Table S6.

**Fig. 2.**
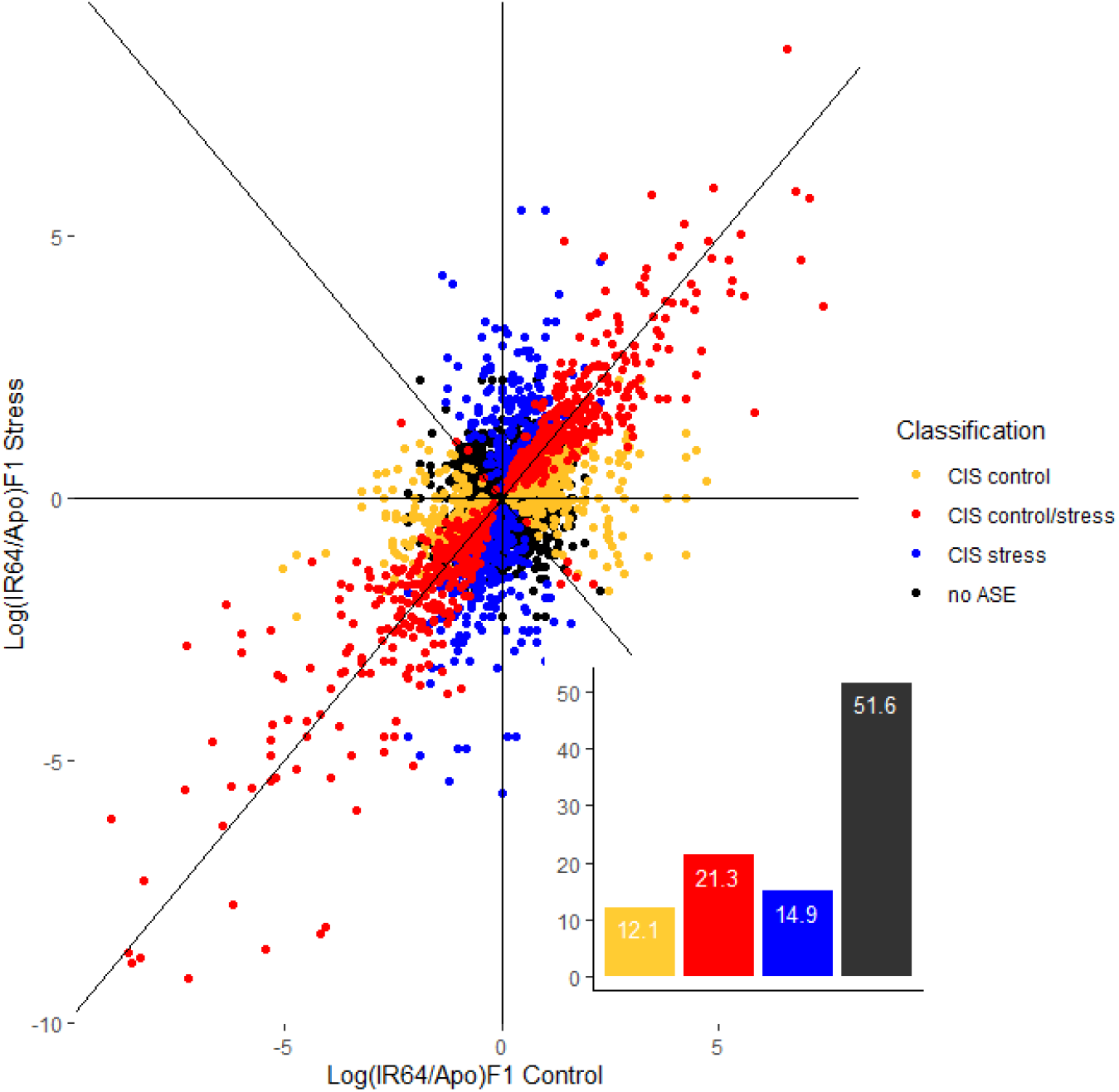
Relative expression ratios between IR64- and Apo-specific alleles in the F1 (Log_2_IR64_F1_/Apo_F1_) under non- (x-axis) and water-stress (y-axis) conditions. Genes are classified based on their CIS category under non- and water-stress conditions (at FDR<0.5%). Inset bar chart: relative proportion (in percentage) of each CIS category including genes with no ASE classification.

Of the 3058 genes that satisfied our criteria (see Materials and Method), 651 isoforms or 21.3% exhibit AI under both conditions (binomial exact test, FDR<0.5%; red dots in Fig. 2); 371 isoforms or 12.1% (yellow dots), under non-stress; 457 or 14.9% (blue), under stress conditions. In summary, almost half of the genes (48.4%) in the hybrids exhibited ASE imbalance across the two conditions (at FDR<0.5%). However, these proportions are influenced by FDR levels. 1988 of the 3058 (or 65%) are asymmetrically expressed at FDR < 5% (see Fig. S6a and Table S7). Hence, ASE imbalance as a consequence of cis divergence between the two indica genotypes is pervasive between the two closely related genotypes. In this study, asymmetric expression was found to be more prevalent under water-limiting compared to non-stress conditions (in both FDR levels).

IR64-specific allele is favorably expressed in the hybrid over Apo-specific allele under non-stress conditions with 133 and 95 isoforms, respectively, exhibiting ASE imbalance or cis divergence (includes CIS control only; excludes CIS control/stress) (at FC≥2; binomial exact test, FDR<0.5%; Table S6). However, under drought conditions, Apo-specific allele is favorably expressed over IR64-specific allele with 157 and 134 isoforms, respectively, exhibiting cis divergence (includes CIS stress only; excludes CIS control/stress) (at FC≥2; binomial exact test, FDR<0.5%; Table S6). There is an apparent genome-wide switching of preferential expression from IR64- to Apo-specific allele from non- to water-stress conditions, respectively, indicating that the Apo-derived alleles were favored under stress in the hybrid.

Interestingly, 11 genes exhibit allele preference switching in the hybrid, shown as red dots lying closely in the y = –x axis in Fig. 2. These are isoforms which switch from IR64- to Apo-specific allele (or vice-versa) between well-watered and water-stressed conditions. This finding suggests that preferential expression (and possibly dominance) in the heterozygote is condition-dependent. This is an interesting area of further research and needs additional confirmation.

#### Cis-diverging genes co-localize with drought-yield QTL regions

Previous papers indicated that cis-diverging genes play major roles in interspecific variations and adaptive divergence. In the current paper, we co-localize cis-diverging genes with marker regions previously identified to participate in drought–yield response (Ereful et al. 2016; Venuprasad et al. 2009; see Materials and Method). Results suggest that several of these cis-diverging genes are aligning with regions underlying drought–yield QTLs in chromosomes 1, 2, 3, 8 and 12 (Fig. 3).

**Fig. 3.**
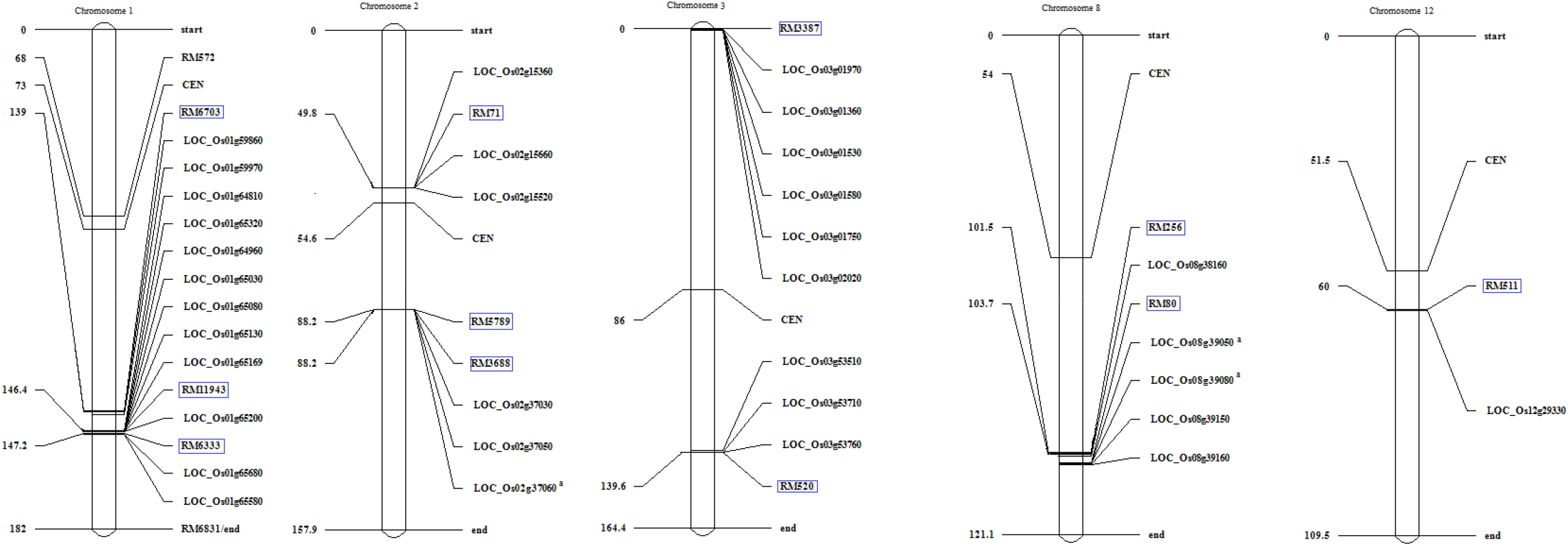
Several genes harboring cis divergence in either or both conditions (binomial exact test, FDR<0.5%) were found to co-localize with QTL marker regions (boxed in blue) identified to participate in drought-yield response in the same bi-parental population background (IR64/Apo) under drought conditions (Venuprasad et al. 2009; Ereful et al. 2016). Markers and genes are said to co-localize if their estimated distance is less than 240kb, the estimated equivalent of 1 cM. (CEN, centromere. Genes appended with a superscript (^a^) were found to harbor cis-divergence at FDR<5% (binomial exact test). Map was generated using ICIMapping (Meng et al. 2015). Positions of markers were estimated based on Cornell and IRMI SSR map at www.archive.grameme.org. Figures not drawn to scale).

Most notably, the drought-response gene LOC_Os12g29330, in particular, tightly co-localizes with RM511, with an estimated distance of 9 kb. It is highly expressed in IR64 as indicated by the positive ratios in both parental (IR64/Apo) and hybrid genotypes (F1_IR64_/F1_Apo_) in both non- and water-stress conditions (Table S2 and S4, respectively). However, while IR64 expresses this TF, it is drought-susceptible. Rice SNP-Seek database (Alexandrov et al. 2015; Mansueto, et al. 2016) and IR64 assembled draft scaffolds (Cold Spring Harbor Lab (CSHL); Schatz et al. 2014) confirm the structural differences of the 2K upstream region of this locus between the two genotypes, i.e. alignments show sequence variations (SNPs and InDels) when pairwise-aligned (Fig. S7; Fig. S8). A multiple intraQTL in the qDTY_12.1_ region centered on the NAM gene (OsNAM_12.1_) was suggested to provide a concerted drought response (Dixit et al. 2015). In our study, all genes in the qDTY_12.1_ region but one (LOC_Os12g29220, nodulin Mt3, OsMtN3_12.1_) co-localizing with the NAM gene were not expressed (Table S1). Thus, the expression of this suite of genes in IR64 through genome editing and other traditional or conventional platforms may be an interesting area of study in the future.

Another set of cis-diverging genes is aggregating in the RM11943–RM6333 region. This was found to be a major QTL (qDTY_1.1_) for grain yield under reproductive-stage drought stress, with a consistent effect in multiple elite lines (Vikram et al. 2011). The most interesting genes on this region include LOC_Os01g64810 (Zinc Finger), LOC_Os01g64960 (chlorophyll a/b binding protein), LOC_Os01g65080 (C2H2–Zinc Finger), and LOC_Os01g64790 (AP2). TFs (such as zinc fingers and AP2) and chlorophyll a/b binding protein have been known to participate in drought tolerance (e.g. Han et al. 2020; Xie et al 2019; Xu et al. 2012).

In chromosome 2, LOC_Os02g15360 (expressed protein), LOC_Os02g15660 (tetratricopeptide repeat), and LOC_Os02g15520 (transposon), which are found to diverge in cis align with RM71. On the other hand, the contiguous genes LOC_Os02g37050 (phosphatidyl inositol glycan), LOC_Os02g37030 (protein binding protein), and LOC_Os02g37060 (photosystem II protein) coincide with the RM5789–RM3688 region.

LOC_Os03g09170 (ethylene responsive TF) and LOC_Os03g02020 (stress responsive A/B barrel domain) are most interesting in chromosome 3 which align with RM3887. Furthermore, LOC_Os03g53510 (WD domain, G-beta repeat domain), LOC_Os03g53760 (ATP-dependent RNA helicase), and LOC_Os03g53710 (aldose 1-epimerase) exhibit cis-divergence and co-localize with RM520.

In chromosome 8, LOC_Os03g58160 (heat stress TF) aligns with RM256; LOC_Os03g53710 (cytokinin-O-glucosyltransferase), LOC_Os08g39050 (pentatricopeptide), LOC_Os08g39080 (tetratricopeptide repeat), and LOC_Os08g39160 (formyl transferase) with RM80.

In summary, GO enrichment analysis, ASE test, differential expression and results of co-localization analysis consistently suggest that cis and/or trans architecture may explain the contrasting response of the two genotypes against drought. Taken all together, these converging evidences underscore the association of cis-diverging regulatory regions with drought response.

In *Saccharomyces paradoxus* (Naranjo et al. 2015) and *Drosophila* (Combs et al. 2018; H. Fraser, pers. comm. Aug. 2019), the identification of trait-associated candidate genes using parents and their hybrids is possible. Therefore, we suspect that it may also be possible to identify contrasting or diverging adaptive loci; hence, candidate genes associated with important traits using hybrid and their parental lines in crops without generating mapping population. However, this needs further confirmation.

Finally, mapping cis and/or trans regulatory differences within and between related taxa has been reported in several organisms especially model organisms but is largely unexamined in crops partly due to its highly computational approach. Here, we provided preliminary results suggesting the possibility of assessing cis and/or trans regulatory differences in an important crop using parent–hybrid expression analysis. Combining such approach with QTL mapping is a powerful strategy. However, such platform is yet to be fully exploited for crop breeding.

## Materials and Methods

### Dry-down experiment

Seeds from both parental and hybrid genotypes were initially grown in petri plates under room temperature. After germination, seedlings were transferred to small pots under controlled conditions inside the phytotron, at the International Rice Research Institute (IRRI). Leaf samples were collected to test hybrid status using SSR markers (RM269, RM 511 and RM 80). Hybrid which exhibited clear polymorphism between the parents using these markers were selected for further experiment. Twenty-four (24) seedlings with confirmed polymorphism were transplanted in large pots.

Plant samples were exposed to either non- and water-stress conditions. For water-stressed samples, the fraction of transpirable soil water (FTSW) dry-down approach was applied as previously described (Cal et al. 2013; Serraj et al. 2014; Sinclair and Ludlow 1986). Water-limiting conditions was implemented by initiating a soil dry-down protocol starting 10 days before heading until the plants reached 0.5 FTSW.

### RNA extraction

At the end of the dry-down treatment, leaf samples from each plant were collected. Samples were snap-frozen in liquid nitrogen and stored at -80°C. RNA was extracted using the TRIzol method according to the instructions provided by the supplier (Invitrogen, San Diego, Calif., USA). RNA-seq libraries were prepared as described in Illumina’s standard protocol for RNA-seq using the parental (IR64 and Apo) and F1 RNA samples from each treatment (non- and water-stress).

### Bioinformatics pipeline

Quality checking. Libraries were sequenced on Illumina GAIIx, generating 38- and 90-base paired end (PE) reads, for the first and second sequencing protocols, respectively. Each sequencing protocol also corresponds to first and second biological replicate.

Quality checking was performed using FASTQC; trimming using FASTX-Toolkit (http://hannonlab.cshl.edu/). We checked the 75% percentile of the sequencing qualities for each base of the PE reads in each sample replicate. Bases with 75% percentile of the sequencing qualities□<□28 were removed for all reads in the replicate.

Note: Analysis pipeline is shown in S2.

### Creating a consensus pseudo-reference

Our bioinformatics approach of estimating ASE levels in the hybrid employs the construction of an *in silico* transcriptome pseudo-reference sequence. This strategy is similar to Shen et al. (2012) and overcame read-mapping biases which may distort ASE estimation (Degner et al. 2009). Use of reference-base mapping (i.e. Nipponbare MSU v7) to align the heterozygote may cause bias (Degner et al. 2009); that is, higher mapping rates towards the allele in the reference sequence. Thus, we preferred the creation of a pseudo-molecule or an *in silico* transcriptome reference to map the reads.

The *O. sativa* ssp. *japonica* (cv. Nipponbare) MSU v7 cDNA (http://rice.plantbiology.msu.edu/) was indexed using bowtie2 (Langmead and Salzberg, 2012). Using the same tool, reads from IR64 and Apo were separately mapped against the indexed reference sequence (no-mixed; no discordant). Alignment options (rdg, rfg, score-min) were set at lenient parameters to allow higher percentage alignment since the materials (IR64 and Apo) are indica and the reference is a japonica.

Using SAMtools (Li et al. 2009), variants (SNPs and Indels) were identified between the MSU7 reference cDNA sequence and the sorted alignment files of both genotypes (commands: sort and mpileup). Custom PERL scripts were created to call SNPs and InDels common to Apo and IR64 sequences (*indica*), but different from the Nipponbare reference sequence. SNP reads must have at least 5× coverage and the SNP proportion must be more than 0.8. For InDels, read coverage must also be at least 5× but InDel proportion should be at least 0.5. We then developed another PERL script which incorporates these variants (SNP and Indels common to IR64 and Apo but different from the MSUv7 cDNA) to modify the reference sequence.

We further mapped 93-11 *indica* RNAseq reads from BGI (http://rise2.genomics.org.cn/) against the pseudo-reference and MSU reference transcriptome sequence and found an increase percentage alignment of 1.09%.

Note: All PERL scripts are available upon request.

### SNP calling

Bowtie2 was used to map Apo, IR64 and the F1 genotypes to the pseudo-reference. SAMtools (options: sort and mpileup) was used to sort and parse the mapping results. To find the SNPs between Apo and IR64 sequences, we created a separate PERL script. The criteria, however, should be SNP reads must have at least 3× read coverage and SNP proportion must be more than 0.8. We used these SNPs to discern which read belongs to which genotype in the F1. These variant calls allow us to distinguish and quantify the two genotype-specific SNP-reads in the heterozygote. Reads with multiple SNPs may cause over-estimation of reads. To avoid such scenario, we used a read-wise approach by counting the number of reads with SNPs regardless of their frequency within a read. A separate custom PERL script was created to carry out this step.

### Gene expression estimation and normalization procedure

We counted the number of reads from the parents and the genotype-specific alleles in the F1 using eXpress (Roberts and Pachter 2013). All statistical analyses in this study were performed using the software R (v 3.5.1 CRAN).

Normalization was performed using DESeq2 (Love et al. 2014). We preferred DESeq2 scaling factor for the normalization procedure as it is, along with Trimmed Means of M-values (Robinson and Oshlack, 2010), more robust to varying library sizes (Dillies et al. 2013). Normalized count data were rounded off to the nearest integer for downstream statistical tests such as binomial exact test.

Average of the normalized read counts of the two replicates for each sample was calculated. Only genes with a total average normalized read count of 20 of the parental samples (IR64 + Apo ≥ 20) were considered for further analysis as recommended (Bell et al. 2013; McManus et al. 2010; Shi et al. 2012). In cases where the isoform has zero read count in either of the parents (IR64 = 0 or Apo = 0) or in either of the parent-specific allele in the hybrid (F1_IR64_=0 or F1_Apo_=0), we adjusted 0 to 1 in order to calculate ratios needing positive integers (e.g. log_2_ transformation). The parental read sum however should be at least 20.

### Cis/trans regulatory differences assignment

Normalized read counts of the parental and the hybrid datasets were analyzed for evidence of regulatory divergence using binomial exact tests followed by FDR analysis (Storey and Tibshirani, 2003) at two significant thresholds of 0.5% and 5% FDR. We did not observe any changes in the general trends between the two FDR levels, e.g. dominance of cis followed by trans under non-stress; and dominance of trans followed by cis under water-stress conditions, both of which were consistent at both FDR levels. The most conservative analysis (i.e. FDR < 0.5%) was finally considered in the Results and Discussions consistent with other papers (e.g. McManus et al. 2010, Bell et al. 2013).

Genes significantly expressed in either parental or hybrid genotypes were further analyzed for trans effects by comparing the genotype-specific mRNA abundance in the parents and in the hybrid samples using Fisher’s exact tests followed by FDR analysis at a significant threshold of 0.5%. Regulatory divergence types are classified according to the categories described below. Such analysis and regulatory divergence assignments are adapted from previous studies (Landry et al. 2007; McManus et al. 2010; Wittkopp et al. 2008; Bell et al. 2013; Combes et al. 2015).

Cis only: significant DE in both P and C. No significant T (FDR_P_ < 0.5%, FDR_C_<0.5%, FDR_T_≥0.5%);

Trans: Significant DE in P and T, but not C (FDR_P_ < 0.5%, FDR_C_ ≥ 0.5%, FDR_T_ <0.5%);

Cis + trans (synergistic): significant DE in P, C and T (FDR_P_ < 0.5%, FDR_C_ < 0.5%, FDR_T_ <0.5%). The signs of the log_2_FC of these genes in the parental and hybrid are the same. Cis- and trans-regulatory differences favor expression of the same alleles;

Cis × trans (antagonistic): significant DE in P, C and T (FDR_P_ < 0.5%, FDR_C_ <0.5%, FDR_T_ <0.5%). The signs of the log_2_FC of these genes in the parental and hybrid are different. Cis and trans differences favor expression of opposite alleles;

Compensatory: Significant DE in C but not P. Significant T (FDR_P_ ≥ 0.5%, FDR_C_ < 0.5%, FDR_T_ < 0.5%);

Ambiguous: statistical test exhibits significant DE in only one of the three tests and has no clear biological interpretations (e.g. FDR_P_ ≥ 0.5%, FDR_C_ ≥ 0.5%, FDR_T_ < 0.5%).

Conserved: No significant DE in P and C. No significant T (FDR_P_ ≥ 0.5%, FDR_C_ ≥ 0.5%, FDR_T_ ≥ 0.5%);

Both “conserved” and “ambiguous” genes were omitted from discussions.

We then performed chi-square test (at P<0.05) in our data count between the two contrasting treatments to determine regulatory categories responding significantly with respect to environmental changes. The analysis was performed using 2-sample test for equality of proportions with Yates’ continuity correction. The number of isoforms in each regulatory category from each condition constitute the first vector; the total number of isoforms in each condition detected, the second vector.

DESeq2 multi-factor design was used to test for GDE and DDE. We used the raw data count of 9141 genes expressed with at least one mapped read across genotype – line – treatment to test for DE (Table S2). Only the parental read counts (not the hybrid) were included for this analysis.

GDE was estimated between the genotypes across the two conditions: (IR64 non-stress, IR64 stressed) vs (Apo non-stress, Apo stressed). Raw read counts were used as the input data count with “IR64” and “Apo” as genotypes. Size factors and dispersions were estimated by DESeq2. On the other hand, differences that arise between the two genotypes under drought conditions relative to non-stress (3-way DE or DDE) were also estimated using the model: (IR64_con, Apo_con) vs (IR64_stress) vs (Apo_stress). Using the function “relevel” of DESeq2, we declared “Apo” and “control (non-stress)” as first level parent and condition, respectively. Results were compared to the genes exhibiting cis/trans regulatory divergence under non- and water-stress conditions, respectively, to find any overlaps.

We used AgriGO (Tian et al. 2017) to perform Gene Ontology (GO) enrichment analysis located at http://systemsbiology.cau.edu.cn/agriGOv2/, implementing Singular Enrichment Analysis (SEA) with *O. sativa* MSU v7 as the reference background.

### Binomial exact test between the two genotype-specific alleles in the hybrids

A binomial exact test between the two parent-specific alleles in the heterozygote under non- and water-stress conditions was performed using R (CRAN v3.4.2). This was followed by FDR analysis at a significant threshold of 0.5%. Such analysis was adapted from Cubillos et al. (2014). An isoform should have a total read count (average normalized read counts of parent-specific alleles) of 20 or more (F1_Apo_ + F1_IR64_ ≥ 20) in one or both conditions. Isoforms with a total read count of 0 in one or both conditions are discarded. Their log-transformed ratios could not be calculated and binomial exact test could not be performed.

### Pairwise-comparison between Apo and IR64 draft genomes

As it is an established drought-tolerance mainstay, the promoter sequence (2K upstream region) of LOC_Os12g29330 (NAM) was studied in this paper. We used the published draft genomes of both Apo and IR64 available from Rice SNP-Seek database and assembled IR64 draft scaffolds. To confirm cis regulatory divergence between the two genotypes, *in silico* promoter sequence analysis was performed. Sequence comparison of the promoter sequences between the genotypes was initially visualized using JBrowse located on SNP-Seek (http://snp-seek.irri.org/jbrowse). As gaps may be ascribed to absence of sequence coverage, both IR64 and Apo datasets were downloaded from the site to confirm their structural variations. We further confirmed whether there are differences between the genotypes by pairwise-aligning genes and their promoter regions from Apo (Rice SNP-Seek database) against IR64 assembled draft scaffolds (CSHL).

BLAST alignments between IR64 (SNP-Seek) and IR64 (CSHL) using several cDNA query sequences show slight disagreement between the two. We preferred the latter reference assembly for our succeeding analysis since IR64 (CSHL) – Apo (SNP-Seek) alignment exhibited more conservative structural variations; hence, higher alignment score than IR64 (SNP-Seek) – Apo (SNP-Seek).

### Co-localization procedure

Co-localization procedure was performed by aligning the positions of cis-diverging genes (Fig. 2; Table S6) to previously identified QTL markers (viz. SSR). These QTL regions were known to be involved with yield under drought in Recombinant Inbred Line (RIL) populations with the same parental background (IR64/Apo): F_3:5_ (Ereful et al. 2016) and F_2:3_ (Venuprasad et al. 2009). These SSR marker regions were anchored in the indica genome and their locations were estimated using Ensembl. Using the same tool, their proximity with genes exhibiting cis-divergence were estimated. Several genes were found to co-localize with the QTL markers at a physical distance of at most 240 kb, the estimated equivalent of 1 cM. The application ICIMapping (Meng et al. 2015) was used to generate the map based on Cornell and IRMI.

## Data Availability

All sequencing data from this work are available at NCBI Sequence Read Archive with a submission entry: SUB1568816 with BioProject ID PRJNA338445.

## Acknowledgement

This work is supported in part by a grant from Biotechnology and Biological Sciences Research Council (BBSRC: BB/F004265/1.), National Institute of Agricultural Botany (NIAB), and the International Rice Research Institute (IRRI). We thank Jack Lagare and Connie Lotho of the IRRI for providing figures of rice pedigree trees. We also thank Patricia Wittkopp and Joe Coolon for additional help in the scripting during the analysis.

## Supplementary materials

S1. Pedigrees of IR64 and Apo, taken from IRIS.

S2. Analysis pipeline for determining regulatory divergence and ASE.

Table S1.List of isoforms and their corresponding raw read counts and average normalized read counts which have a minimum mapped read of one or more for at least one column across genotype–line–treatment combinations

Table S2. Isoforms identified to have at least 20 mapped reads in either of the parental genotypes under non-stress conditions and their regulatory divergence classification

Table S3. Results of GDE showing isoforms genotypically differentially expressed (FC≥2; FDR<0.05)

Table S4. Isoforms identified to have at least 20 mapped reads in either of the parental genotypes under water-stress conditions and their regulatory divergence classification.

Table S5. Results of 3-way differential expression or drought DE (DDE).

Table S6. Results of ASE binomial exact test (FDR<0.5%). Isoforms were classified as cis-control, cis-stress, or cis (control/stress).

Fig. S1. Graphical results of GO enrichment analysis (biological process) using cis-diverging genes under non-stress conditions as input sequence.

Fig. S2. Graphical results of GO enrichment analysis (cellular component) using cis-diverging genes under non-stress conditions as input sequence.

Fig. S3. Graphical results of GO enrichment analysis (biological process) using trans-diverging genes under non-stress conditions as input sequence.

Fig. S4. Graphical results of GO enrichment analysis (cellular component) using trans-diverging genes under non-stress conditions as input sequence.

Fig. S5. Graphical results of GO enrichment analysis (biological process) using trans-diverging genes under water-stress conditions as input sequence.

Fig. S6. Graphical results of GO enrichment analysis (cellular component) using trans-diverging genes under water-stress conditions as input sequence.

Fig. S7. Pairwise-alignment results using BLAST between IR64 (CSHL) and Apo (SNP-seek) of the promoter region for the gene, No apical meristem (LOC_Os12g29330)

Fig. S8. Graphical representation of the pairwise alignment of the promoter sequence for the gene, No apical meristem (LOC_Os12g29330) using SNP-seek genome browser.

