## Supplementary figures and images for "cis dominantly explains regulatory divergence between two indica rice genotypes; drought further enhances regulatory differences"

### Fig. S1

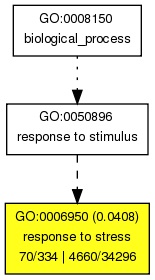

### Fig. S2

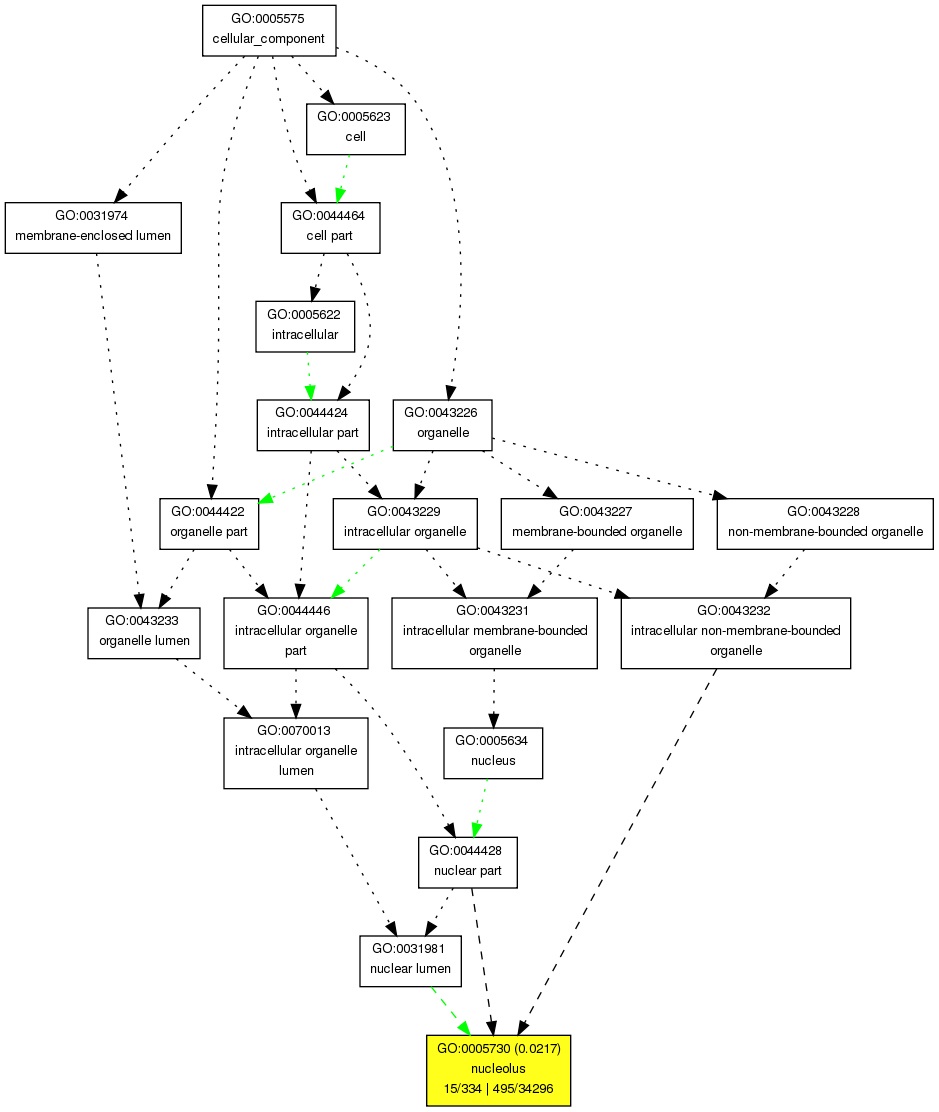

### Fig. S3

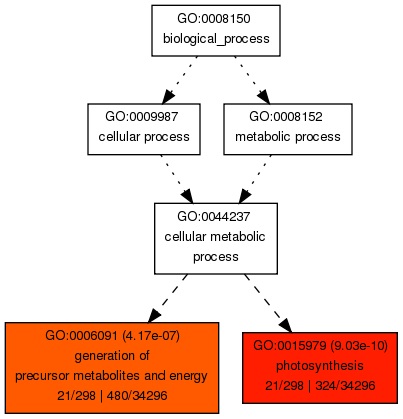

### Fig. S4

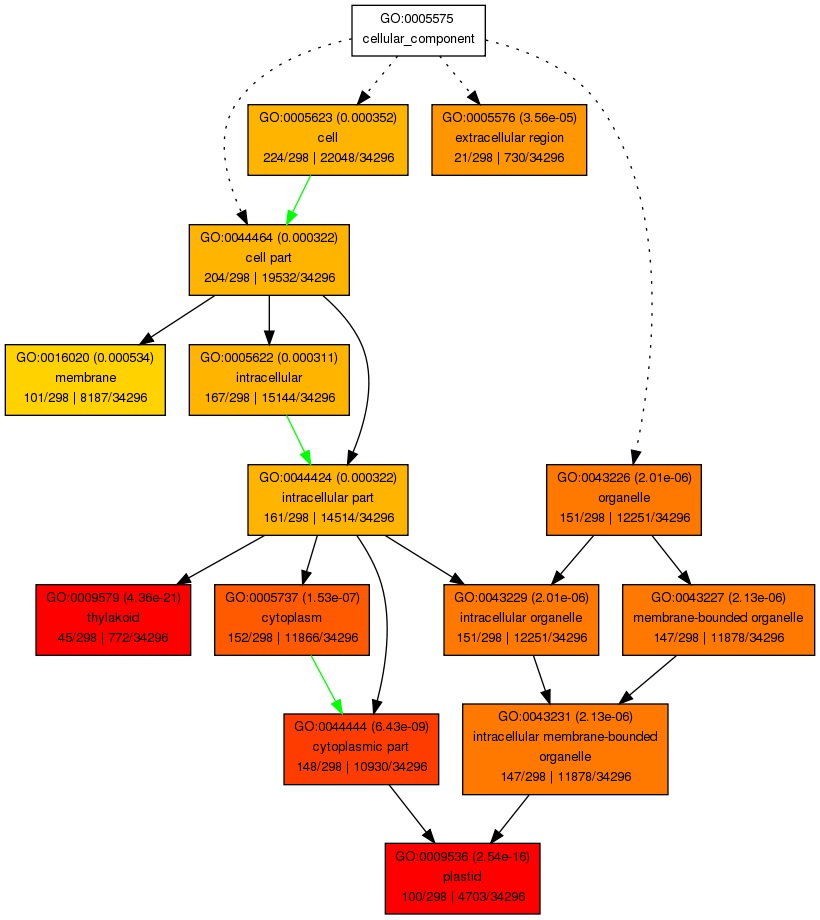

### Fig. S5

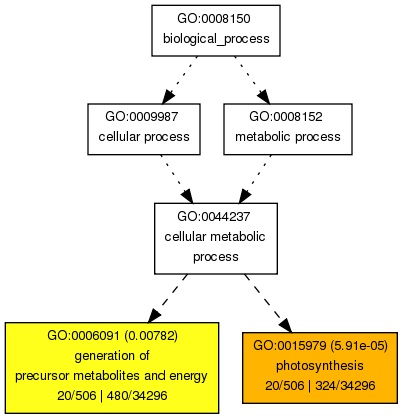

### Fig. S6

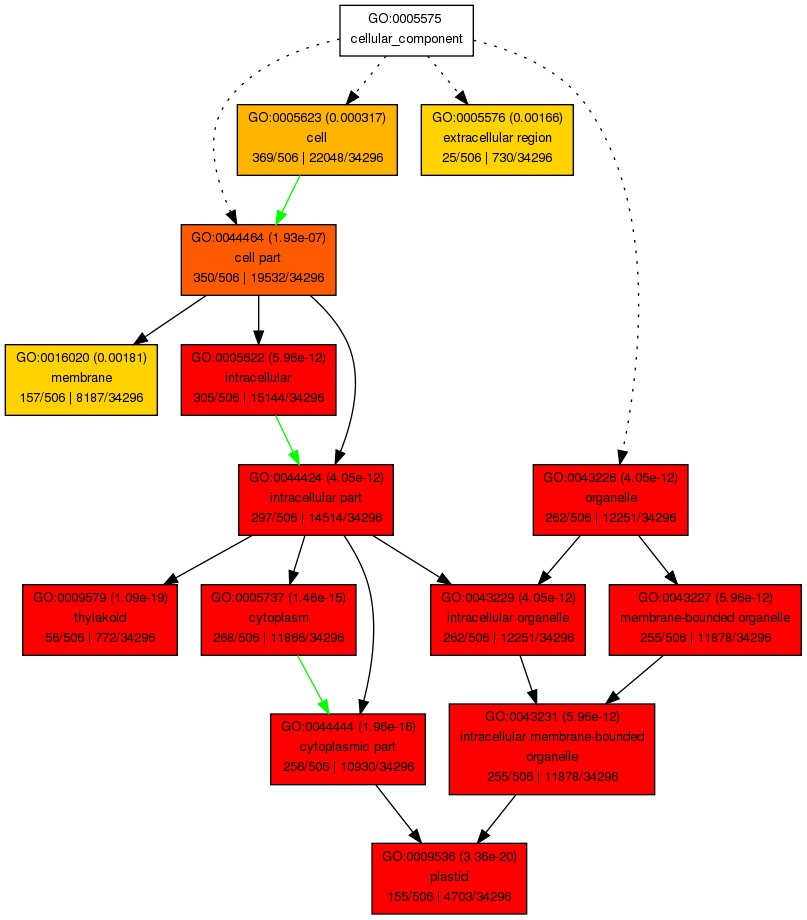

### Fig. S8

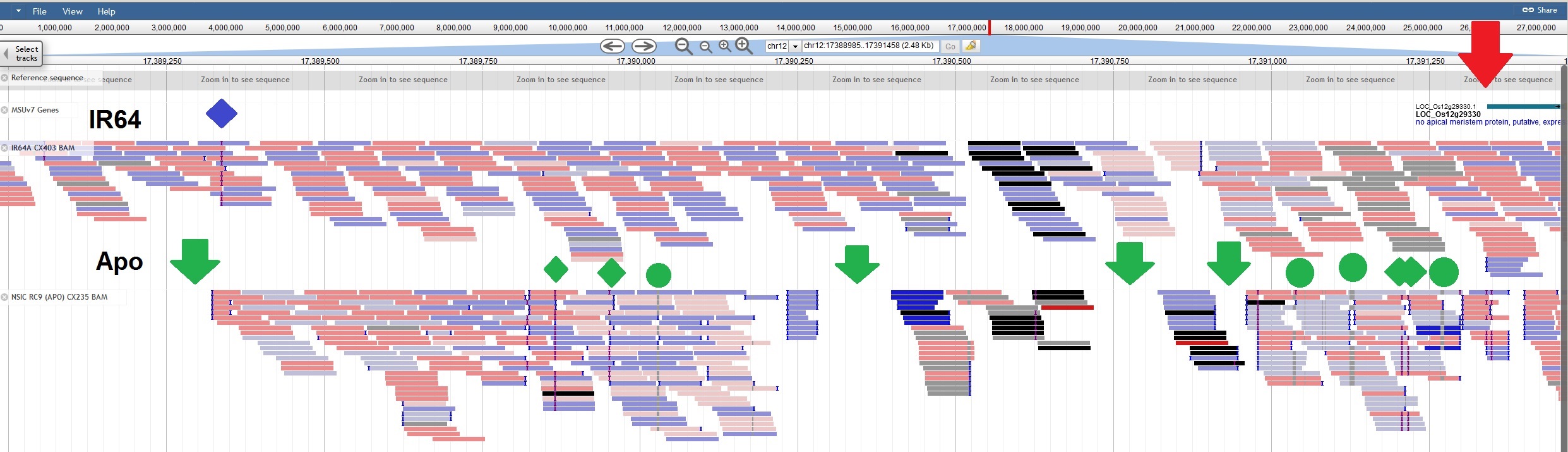
