## Supplementary material for "cis dominantly explains regulatory divergence between two indica rice genotypes; drought further enhances regulatory differences": S1

```
Pedigree tree for IR 64

PNO   NAME AND LEVEL                                              DSTEP TERM GID
1    IR 64                                                            3    50533
5    +-IR 5657-33-2-1                                                 3    24486
10   | +-IR 5236                                                      0    8180
18   | | +-IR 2006-P3-31-3                                            3    8082
33   | | | +-IR 1870                                                  0    3299
62   | | | | +-IR 1103-15-8-4-2-2-3                                   6  < 3290
63   | | | | +-IR 1163-124-1-3                                        3  < 3298
34   | | | +-IR 1614                                                  0    2723
64   | | |   +-IR 22                                                  2  < 2722
65   | | |   +-IR 1402                                                0  < 2201
19   | | +-IR 2146-68-1                                               2    8111
37   | |   +-IR 773 A 1-36-2-1                                        4    3411
70   | |   | +-IR 8                                                   2  < 715
71   | |   | +-IR 238-28-3-2                                          3  < 1127
38   | |   +-IR 1916                                                  0    3414
72   | |     +-IR 773 A 1-36-2-1                                      4  * 3411
73   | |     +-IR 1915 B                                              0  < 3413
11   | +-IR 5338                                                      0    8307
20   |   +-IR 2061-465-1-4                                            3  * 8301
16   |   | +-IR 833-6-2-1-1                                           4    3707
26   |   | | +-IR 262-43-8-11                                         3  < 1171
27   |   | | +-GAM PAI 30-12-15                                       3    109
55   |   | |  >GAM PAI                                                ?  # 253220
17   |   | +-IR 2040                                                  0    3661
28   |   |   +-IR 1561-149-1                                          2  < 3660
29   |   |   +-IR 1737                                                0  < 3006
21   |   +-IR 2055-475-2                                              2    7326
46   |     +-BPI 121-407                                              ?    3680
78   |     | +-FORTUNA                                                1    421581
135  |     | +-PA CHIAM                                               ?  # 420345
79   |     | +-SERAUP BESAR 15                                        ?  # 320386
47   |     +-IR 1833                                                  0    3199
80   |       +-IR 1416-131-5                                          2  < 3161
81   |       +-IR 22                                                  2  < 2722
6    +-IR 2061-465-1-5-5                                              4    11072
16     +-IR 833-6-2-1-1                                               4    3707
26     | +-IR 262-43-8-11                                             3    1171
51     | | +-PETA                                                     ?    11
83     | | | +-CINA                                                   ?  # 168602
84     | | | +-LATISAIL                                               1    168603
140    | | |  >LATISAIL                                               ?  # 2271264
52     | | +-IR 95                                                    0    155
85     | |   +-PETA                                                   ?  * 11
86     | |   +-IR 86                                                  0  < 138
27     | +-GAM PAI 30-12-15                                           3    109
55     |  >GAM PAI                                                    ?  # 253220
17     +-IR 2040                                                      0    3661
28       +-IR 1561-149-1                                              2    3660
58       | +-IR 579-48-1-2 (NILO 11)                                  3    2583
90       | | +-IR 8                                                   2  < 715
91       | | +-TADUKAN                                                2    815
144      | |  >TADUKAN                                                ?  # 2271990
59       | +-IR 747-B2-6-3                                            4    2577
96       |   +-TKM 6                                                  ?  < 441
97       |   +-IR 746 A                                               0  < 1076
29       +-IR 1737                                                    0    3006
60         +-IR 24                                                    3    2444
101        | +-IR 8                                                   2  < 715
102        | +-IR 127-2-2                                             2  < 960
61         +-IR 1721                                                  0    2983
103          +-IR 24                                                  3  * 2444
104          +-IR 1704                                                0  < 2933
 NAME - ">" indicates germplasm derived from germplasm in the line above
        "+" indicates generative progenitor of the germplasm above the "+"
            female progenitors are above male progenitors
 DSTEP - Number of derivative steps from generative source
 TERM  - Termination code:
         "*" indicates lines expanded elsewhere in the tree
         "<" indicates lines which could be further expanded
         "#" indicates germplasm with unknown parents


PEIGREE TREE for APO

PNO   NAME AND LEVEL                                              DSTEP TERM GID
1    IR 55423-01 (NSIC RC 9)                                          2    204538
4    +-UPL RI 5                                                       ?    406626
8    | +-SIGADIS (AICRIP)                                             ?    386533
17   | | +-BLUEBONNET                                                 ?    96089
33   | | | +-REXORO                                                   2    38864
50   | | | +-MARONG PAROC                                             ?  # 420278
34   | | | +-FORTUNA                                                  2    482
52   | | |  >PA CHIAM                                                 ?  # 420345
18   | | +-BENONG                                                     ?    16717
35   | |  >BENONG                                                     ?  # 2891665
9    | +-BPI 76-1                                                     ?    227416
23   |   +-FORTUNA                                                    1    421581
36   |   +-PA CHIAM                                                   ?  # 420345
24   |   +-SERAUP BESAR 15                                            ?  # 320386
5    +-IR 12979-24-1 (BROWN)                                          3    67816
13     +-IR 10125                                                     0    14281
25     | +-IR 8838                                                    0    12756
37     | | +-IR 3179-25                                               2    12242
55     | | | +-IR 1721-11-8-3-2-3                                     5  < 5495
56     | | | +-C 4-63                                                 ?  < 3695
38     | | +-K PATONG                                                 2    9175
58     | |  >K PATONG                                                 ?  # 33206
26     | +-IR 26                                                      2    7845
41     |   +-IR 24                                                    3    2444
62     |   | +-IR 8                                                   2  * 715
63     |   | +-IR 127-2-2                                             2  < 960
42     |   +-TKM 6 (ACC 237)                                          ?    420745
67     |     +-CO 18                                                  2    421846
93     |     +-VELLAIKAR                                              ?  # 421187
68     |     +-GEB 24                                                 1  < 385404
14     +-IR 879-314-2                                                 2    8771
29       +-IR 8                                                       2    715
45       | +-PETA                                                     ?    11
70       | | +-CINA                                                   ?  # 168602
71       | | +-LATISAIL                                               1    168603
96       | |  >LATISAIL                                               ?  # 2271264
46       | +-DEE GEO WOO GEN                                          2    2
73       |  >DEE GEO WOO GEN                                          ?  # 2285932
30       +-IR 759                                                     0    1089
47         +-IR 8                                                     2  * 715
45         | +-PETA                                                   ?  < 11
46         | +-DEE GEO WOO GEN                                        2    2
73         |  >DEE GEO WOO GEN                                        ?  # 2285932
48         +-IR 407                                                   0    585
74           +-PETA                                                   ?  * 11
75           +-IR 283                                                 0  < 392
 NAME - ">" indicates germplasm derived from germplasm in the line above
        "+" indicates generative progenitor of the germplasm above the "+"
            female progenitors are above male progenitors
 DSTEP - Number of derivative steps from generative source
 TERM  - Termination code:
         "*" indicates lines expanded elsewhere in the tree
         "<" indicates lines which could be further expanded
         "#" indicates germplasm with unknown parents
```
